# Targeting hotspots to reduce transmission of malaria in Senegal: modeling of the effects of human mobility

**DOI:** 10.1101/403626

**Authors:** Kankoé Sallah, Roch Giorgi, El Hadj Ba, Martine Piarroux, Renaud Piarroux, Karolina Griffiths, Badara Cisse, Jean Gaudart

## Abstract

**Background:** In central Senegal malaria incidences have declined in recent years in response to scaling-up of control measures, but now remains stable, making elimination improbable. Additional control measures are needed to reduce transmission.

**Methods:** By using a meta-population mathematical model, we evaluated chemotherapy interventions targeting stable malaria hotspots, using a differential equation framework and incorporating human mobility, and fitted to weekly malaria incidences from 45 villages, over 5 years. Three simulated approaches for selecting intervention targets were compared: a) villages with at least one malaria case during the low transmission season of the previous year; b) villages ranked highest in terms of incidence during the high transmission season of the previous year; c) villages ranked based on the degree of connectivity with adjacent populations.

**Results:** Our mathematical modeling, taking into account human mobility, showed that the intervention strategies targeting hotspots should be effective in reducing malaria incidence in both treated and untreated areas.

**Conclusions:** Mathematical simulations showed that targeted interventions allow increasing malaria elimination potential.

## Background

Malaria remains a major health burden, with a global annual incidence of 212 million new cases and 429,000 deaths in 2015, most of which (92%) occurred in sub-Saharan Africa and mostly (70%) in children under five years ^1^. In line with the situation in Senegal nationwide, malaria incidence has declined in the Mbour area since the 2000s, due to scaling-up of malaria control. This is primarily due to universal coverage of long-lasting insecticide-treated bednets (LLIN) ^2^, improved access to diagnosis (Rapid Diagnostic Tests RDT) and prompt treatment of malaria with Artemisinin-based Combination Therapy (ACT) ^3,4^. Senegal is still in the control phase of the malaria program, according to the World Health Organization (WHO) classification (more than 5 cases per 1,000 inhabitants per year), but the country is committed to achieving the objectives of pre-elimination stage by 2020 ^5^.

Malaria control and elimination projections are challenging due to the complex interactions between humans, vectors, parasite genetic complexity, environmental and socioeconomic factors. Spatial heterogeneity of incidences characterizes low-transmission settings within non-endemic areas of sub-Saharan Africa and Asia ^6,7^. Hotspots are often broadly defined as areas where malaria transmission exceeds an average level ^8,9^. Targeting interventions to specific hotspots may be efficient in reducing malaria burden in the entire area ^8,10,11^. Operational definitions of hotspots allow the evaluation of the impact of targeted interventions in dry or rainy seasons. Intervention strategies analyzed in this study were:

- Focused Mass Drug Administration (MDA) consists of systematically treating individuals in a selected geographic area with antimalarial drugs, without screening for infection.
- Focused Mass Screen and Treat (MSAT) consists of malaria screening using a rapid diagnostic test and providing treatment to those with a positive test result, in a selected area.
- Seasonal Malaria Chemoprevention (SMC) consists of intermittently administrating preventive antimalarial treatment to children during the main transmission period.
- Long-Lasting Insecticide-treated Nets (LLIN) intend to avoid mosquito bites relying on physical and chemical barriers of manufactured nets.

Human mobility plays a critical role in malaria elimination strategies, leading to reintroduction and resurgence of malaria in treated areas, hampering malaria elimination efforts ^12^.

This study aimed to model the impact of spatially targeted malaria interventions, based on different target definitions, taking into account human mobility and using a meta-population mathematical model based on a Susceptible-Exposed-Infected-Recovered (SEIR) framework, with spatially separated populations that interact with each other via moving individuals.

## Methods

### Data

The dataset analyzed during the current study is available as additional file.

The population data came from 45 villages in the health district of Mbour, Senegal (Figure 1) and were collected from 2008 to 2012 through a health and demographic surveillance system established in central Senegal ^13^. Malaria cases at health facilities were confirmed by rapid diagnostic test and geographical coordinates village centroids recorded using GPS (Global Positioning System) devices. Estimates of rainfall were extracted from Goddard Earth Sciences Data and Information Services Center (http://disc2.nascom.nasa.gov/). The model was implemented using R 3.1.2 free software ^14^ (The R Foundation for Statistical Computing), using {deSolve} ^15^ and {FME} ^16^ packages for numerical solution of differential equations describing transmission. The {Geosphere} package ^17^ was used to estimate Euclidian distances between villages. Graphics were edited with Paint.NET freeware raster graphics editor (dotPDN LLC, Rick Brewster).

**Figure 1.**
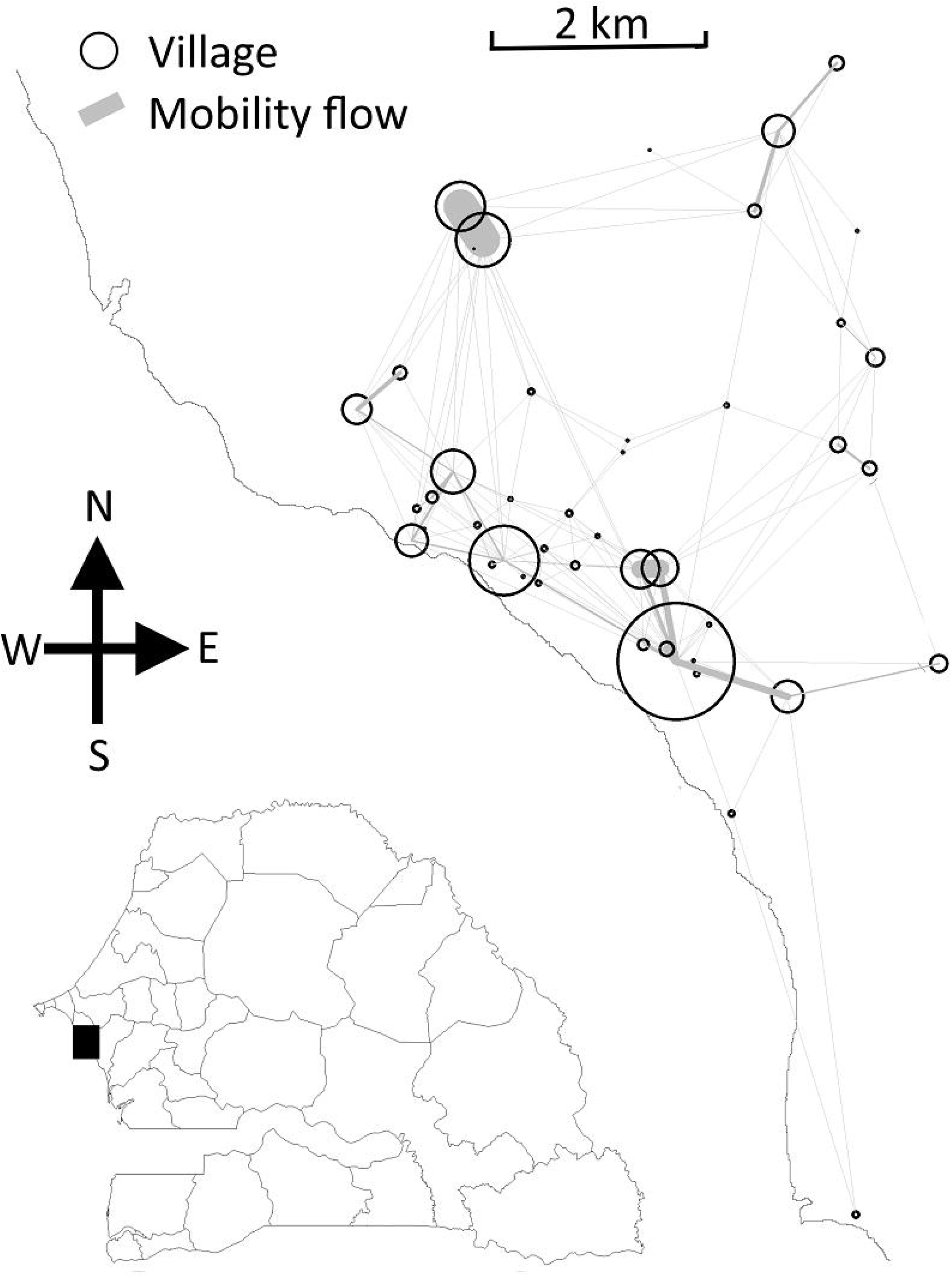
The Mbour zone, Senegal, 2008-2012. Geographical coordinates of the 45 villages are represented by black circles and mobility flows by gray lines. The thickness of the lines reflects the number of trips.

### Model structure

Malaria transmission in each village was represented by a deterministic compartmental SEIR transmission model based on the “Bancoumana” model described by Gaudart et al. ^18^ (Figure 2) Infection removed susceptible individuals from the susceptible compartment *S*_*k*_ (*t*). The proportion of human infection in village *k,* denoted *I*_*k*_ (*t*) was proportional to anopheles density *υ*(*t*), to frequency of mosquito bites *α*, to human susceptibility to infection *β*, and to the effective proportion of infected mosquitoes *i*(*t*). The latter was described as the weighted sum of local proportion of infected mosquitoes *Ai*_*k*_ (*t*) and remoted proportion of infected mosquitoes *Ai*_*j*_ (*t*) (equation 1). The weights depended on the proportion of travelling people at a given time (*m*) and also on relative probabilities *Q*_*kj*_ of travel from remote locations *j* to local village *k*.

**Figure 2.**
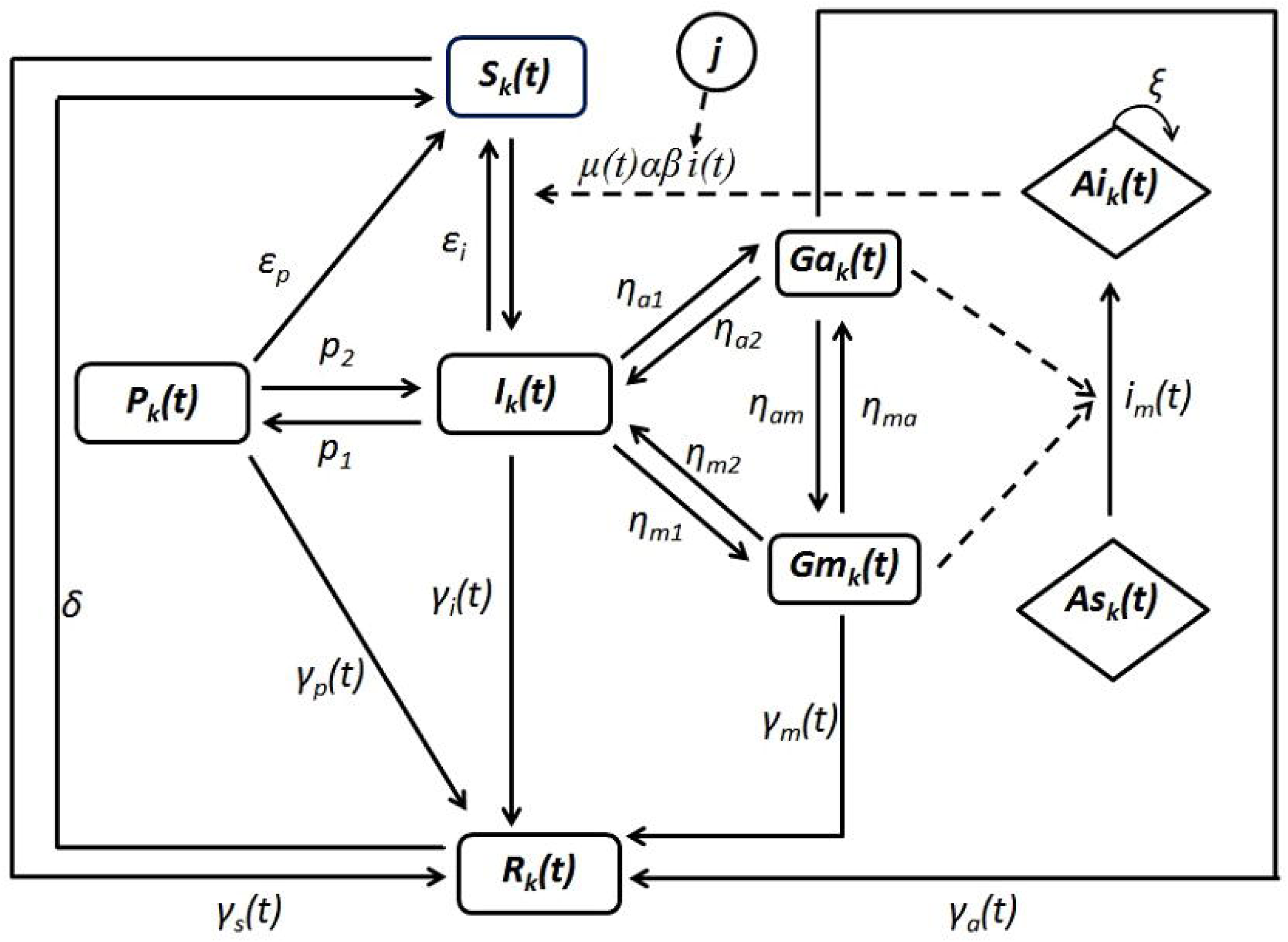
Malaria transmission diagram at a local village *k*. Letter *j* stands for remote villages. Human compartments are *S*_*k*_: susceptible, *P*_*k*_: premunition, *I*_*k*_: blood stage infection, *G*_*ak*_: asymptomatic carriage of gametocytes, *G*_*mk*_: symptomatic carriage of gametocytes, *R*: resistance due to treatment. Mosquito compartments are *A*_*ik*_: infected mosquitoes and *A*_*sk*_: susceptibles mosquitoes. The arrows represent the transition rates between compartments. Other parameters are described in Supplementary information, S1 and S2.

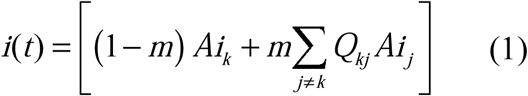

Probabilities *Q*_*kj*_ were estimated via the radiation model of human mobility^19^:

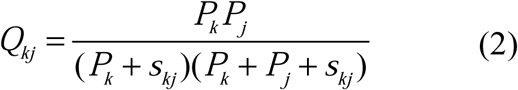

*P*_*k*_ and *P*_*j*_ were the populations sizes in location *k* and *j* respectively and *skj* the total population inside the circle centered at *k* whose circumference touches *j*, excluding the source and destination populations. Travel was modeled as round trips of less than one week duration. Villages populations sizes were assumed constant. Each susceptible inhabitant of a village *k* could infect or be infected at other villages *j*. In this approach, individuals remained residents of their home village but could spend some time in neighboring villages. Long-term mobility was not modeled.

The model assumed that newly-infected individuals *I*_*k*_ (*t*) initially carry only blood stage parasites. Gametocytes subsequently appeared, and the individual may have malaria symptoms or be asymptomatic, leading to respective compartments *Gm*_*k*_ (symptomatic, infectious) and *Ga*_*k*_ (asymptomatic, infectious). All gametocyte carriers were assumed to contribute to transmission. Infection of mosquitoes depended on the effective proportion of human infection *i*_*m*_(*t*), represented as the weighted sum of human infection proportions in the local village and neighboring villages. Gametocytes were transmitted to anopheles by gametocyte carriers resident in the local village *k*, and by gametocyte carriers travelling from other villages *j* to the local village *k*. It was assumed that mosquitoes were infected by feeding on human gametocyte carriers near to mosquito breeding sites, and that blood-fed anopheles did not move from one village to another ^20^.

The model assumed that individuals gradually acquired partial immunity (premunition) at a Rate *p*_1_^21^, after several malaria attacks. Premunition was assumed to be lost at rate *p*_2_. Targeted interventions were modeled as a transition to the compartment of protected, the transition rates being defined as rectangular pulse functions reflecting the administration of interventions over a limited period. Protection resulting from drug administration was assumed to be lost at constant rate.

Seasonal variations of anopheles density *υ*(*t*) were modeled assuming that anopheles density was proportional to the cumulative rainfall over the previous six weeks, oscillating between the minimum and maximum values reported in previous entomological studies within the studied area (0 to 12 anopheles/individual/day) ^22^. The lag between anopheles density and malaria incidences was identified by sensitivity analysis (Supplementary Fig. S1) and the optimal value was about 6 weeks. This value was also consistent with previous studies^23,24^.

The equations of the model are set out in Supplementary Information S1-S2 and a fine description of the parameters or values used in the simulations is shown in Supplementary Table S1.

### Model calibration

The meta-population model was fitted to weekly malaria incidence data from January 1, 2008 to December 31, 2008, using an optimization approach based on Markov Chain Monte Carlo (MCMC) sampling and exploring the probability density function of differential equations parameters ^25^.

Initial values of model compartments were defined as conditions values at the beginning of each rainy season. The average dry season malaria incidence in each village was computed from all the available data and used as the initial value for symptomatic malaria compartment (*Gm*). The proportion of people in the premunition compartment at the start of the simulation (proportion of asymptomatic carriage) relied on expert advice and values from the literature ^26^ (Supplementary Table S2).

The model was validated against data from 2010 to 2012 (See Supplementary Fig. S2. for calibration and validation results). The sensitivity of model parameters was assessed by varying them around the estimated value.

### Hotspots definitions and interventions

Three deliberately pragmatic definitions of hotspot villages were investigated:

1.Low Transmission Period Hotspots (LT Hotspots) were defined as villages reporting at least one malaria case during the low transmission period (December to May) of the previous year.

2.High Transmission Period Hotspots (HT hostpots) were villages with the highest malaria incidences during the last transmission season (June to November).

3.High Connectivity Hotspots (HC hotspots) were villages highly connected to neighboring villages on the basis of human mobility potential.

Human mobility potential was approximated by a degree centrality score (equation 3). The degree centrality score of a village *k* (*d*_*k*_ ) was defined as the number of travel connections from remote villages to the village *k* which amount was above the first decile of the total amount of travels towards *k* ^27^. The degree centrality score helped to capture infection routes from remote villages, and its higher values indicated an increased vulnerability to malaria spread.

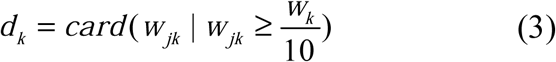

In equation 3, *d*_*k*_ represented the degree centrality score of village *k, card* (cardinality) represented the number of connections from remote villages to village *k* above the threshold 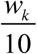, *w*_*jk*_ the number of trips from villages *j* to village *k*, and *w*_*k*_ the total amount of travels towards village *k.*

These hotspot definitions were chosen as they are simple to apply in practice and therefore more practical than definitions that have been based on more formal spatial analysis or requiring serological surveys ^8,10,28^.

Interventions were simulated from 2010. MSAT and MDA drug interventions assumed the use of Dihydroartemisinin plus Primaquine. Coverage was assumed to be 70% for each round of MDA/MSAT, meaning that 70% of the population in targeted hotspots effectively received the intervention (treatment, in the case of MDA and screening then treatment for MSAT). Two rounds of intervention, separated by one month interval, were assumed for both MDA and MSAT. Drugs were provided during the first week of September and again during the first week in October (referred to as HTP: High Transmission Period), or in February and March (referred to as LTP: Low Transmission Period).

The SMC strategy targeted only children under 10 years old, assumed to represent 30% of the population ^29^. Delivery occurred on the first 4 days of each month from September to December, in the whole area under study (all 45 villages). According to WHO recommendations, SMC should not represent a hotspot targeted strategy.

For the sake of simplification, the impact of long-lasting insecticidal nets was implemented as a direct decrease in the rate of mosquito bites (α) over the intervention period.

The intervention efficacy, Δ_*I*_ was defined as the relative variation in malaria annual incidence from no intervention assumption to intervention assumption.

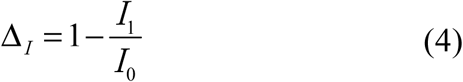

*I*_*0*_ and *I*_*I*_ were annual cumulate malaria incidences respectively before and after intervention.

## Results

### Parameters estimates and sensitivity analysis

The estimated weekly mobility rate was m = 0.09 (95% confidence interval (CI): 0.0015, 0.2) corresponding to 2-200 individuals moving between villages per 1,000 inhabitants per week. The entomological inoculation rate (EIR) calculated from the model, varied seasonally between 0 and 2.16 infected bites per person per night.

Key parameters were varied to assess their sensitivity on malaria incidences (Figure 3). Model predictions were sensitive to the following parameters: anopheles density (33% increase in malaria incidence while increasing parameter by about 5%), access to treatment (16% increase in malaria incidence while decreasing parameter by about 5%), loss of premunition (4.5% increase in malaria incidence for 5% parameter increase) and human mobility (1% increase in malaria incidence for 100% parameter increase).

**Figure 3.**
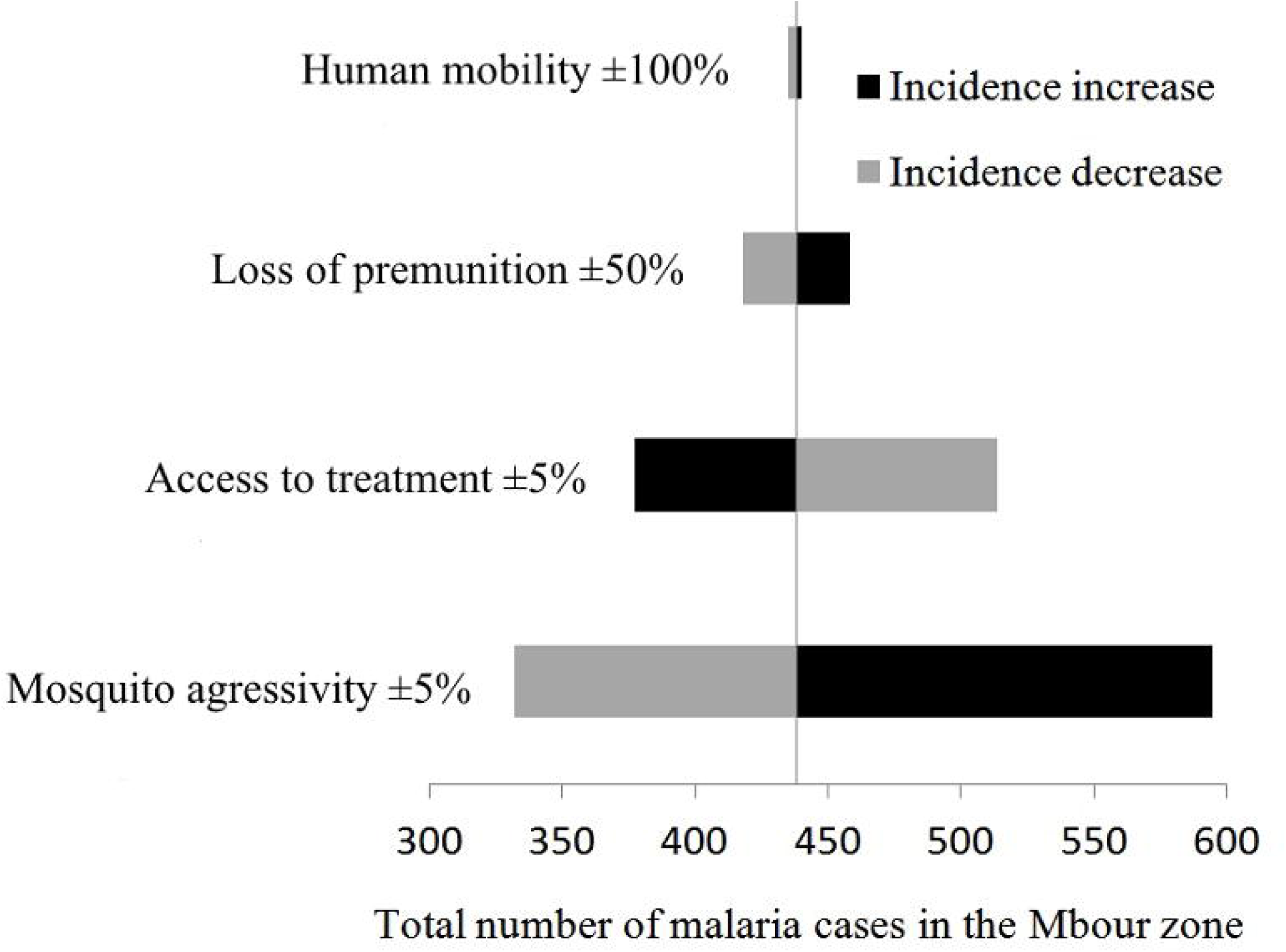
Sensitivity of model parameters in the malaria meta-population model, Mbour, Senegal 2008-2012. Right and left correspond to parameter increase or decrease respectively (percentages). Black and gray bars represent respectively increase and decrease in total malaria cases, subsequent to parameter variations.

### Sensitivity of hotspot definitions

LT hotspots showed temporal instability. Their locations changed from one year to another (Cohen’s Kappa coefficient 0.21, 95% CI: (0.16, 0.33) versus 0.6, 95% CI: (0.36, 0.85) for HT hotspots). HC hotspots were static in time because the definition relied on radiation model mobility flow estimates based on population densities. Annual malaria incidence rates steadily increased in LT hotspots villages compared to other villages, suggesting that LT hotspots might display a qualitative illustration of the quantitative approach defining HT hotspots.

HT hotspots were less populated than LT hotspots (average population per hotspot, 510 inhabitants versus 1,703 inhabitants, Wilcoxon test *P*=0.13), demonstrating that small villages had higher incidence rates during the transmission season.

HC hotspots were slightly more populated than LT hotspots (average population per hotspot, 1,876 inhabitants versus 1,703 inhabitants, Wilcoxon test *P*=0.6) and demonstrated lower malaria incidences than LT hotspots (Wilcoxon test *P*=0.03).

### Intervention simulations

Short and long term variations in annual incidences after a unique intervention and after yearly repeated interventions on LT hotspots are shown on Figure 5, for the overall zone. For HT hotspots and HC hotspots, see Supplementary Fig. S3 and S4, Supplementary Tables S3.a, S3.b, S3.c and S3.d. The simulations performed assuming higher spatial heterogeneity in anopheles density showed no substantial differences between strategies (Supplementary Fig. S5).

**Figure 4.**
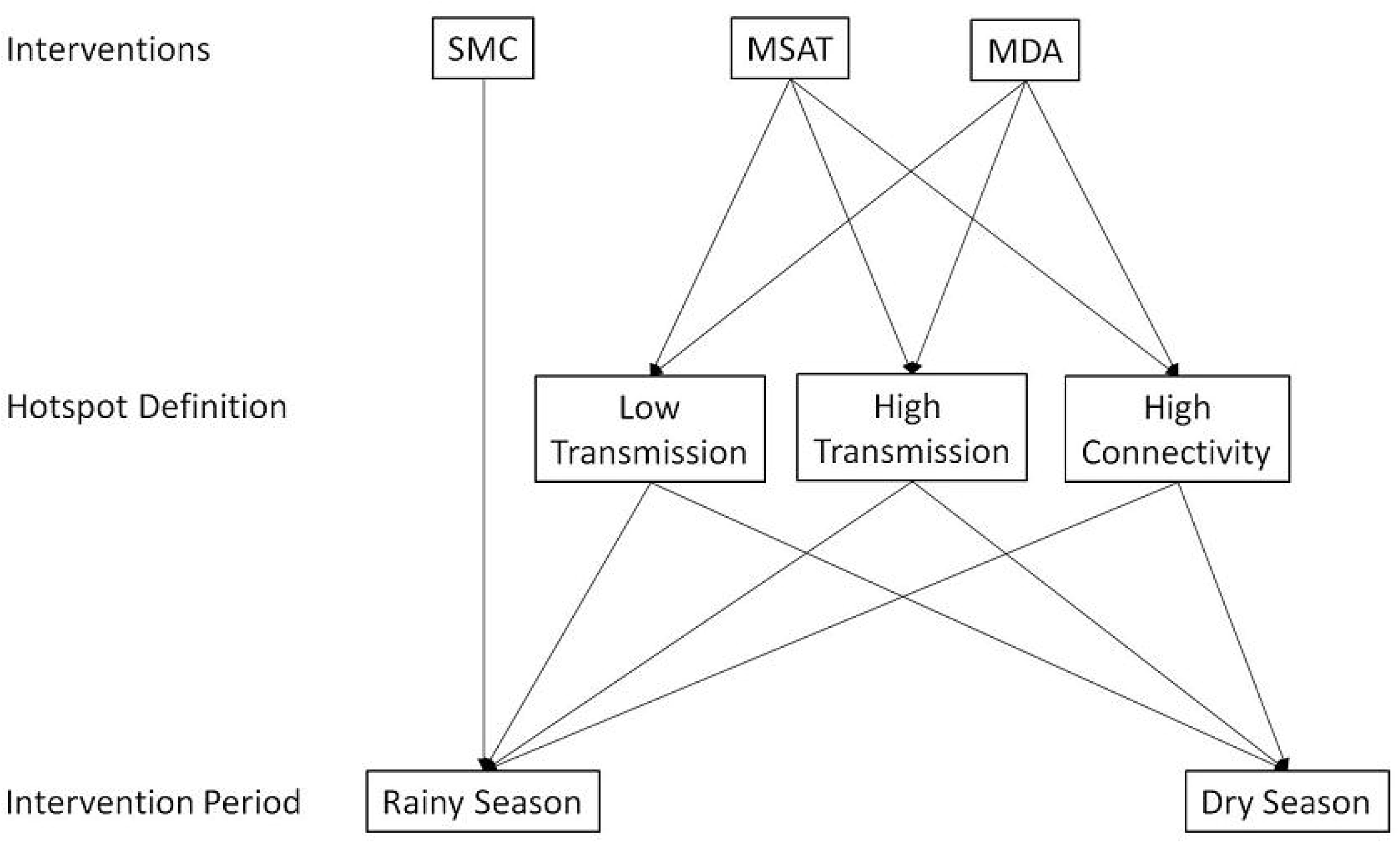
Malaria drug interventions, targets and periods. SMC is recommended to be implemented at beginning of rainy seasons whether in a hotspot or not.

**Figure 5.**
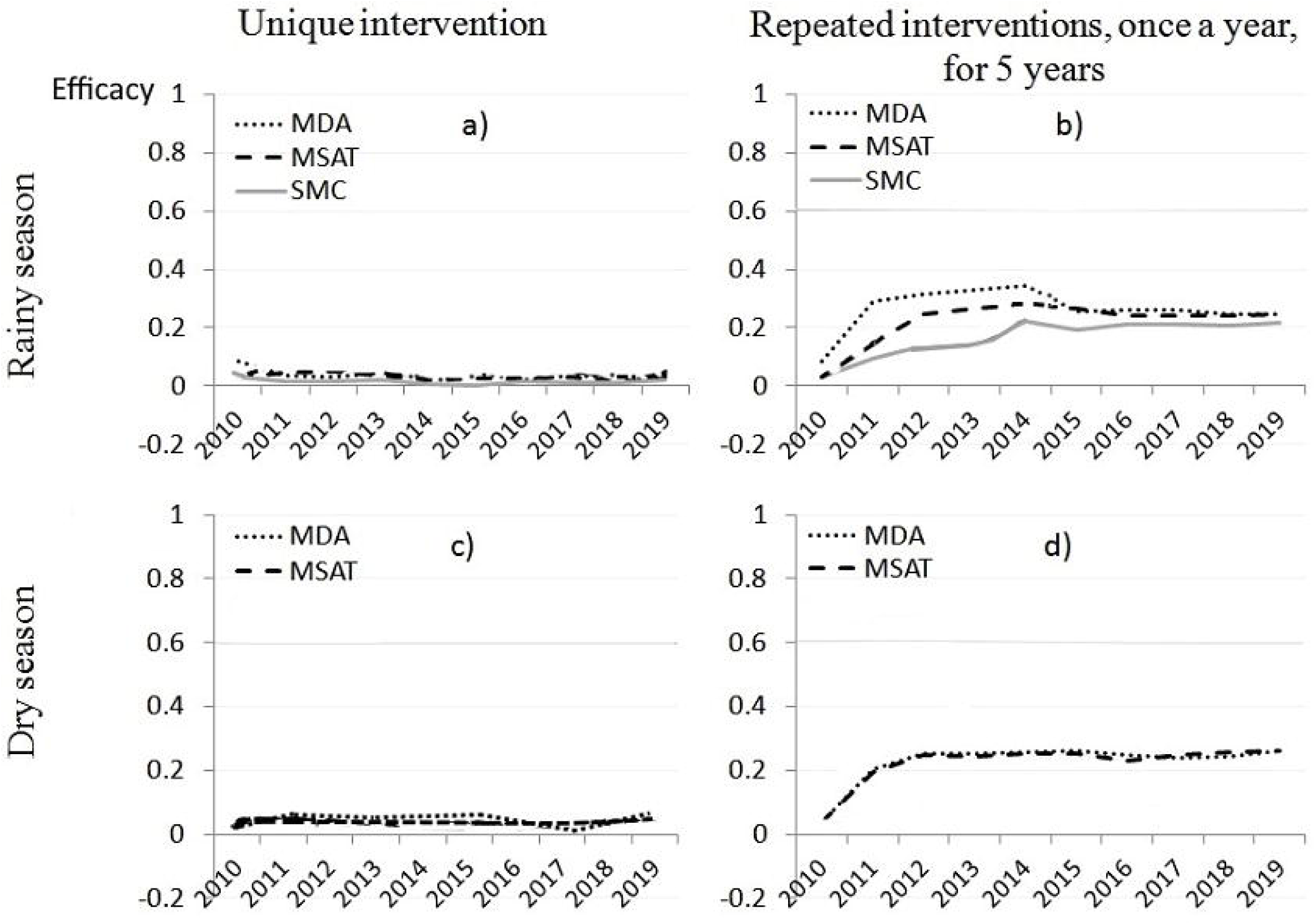
Decrease in malaria incidence while targeting Low Transmission period hotspots (LT hotspots), Mbour, Senegal 2008-2012. Y-axis represents the percentage of decrease in malaria incidence for the overall area (45 villages). a) Unique one year intervention in rainy season, b) Repeated interventions over five consecutive rainy seasons, once a year, c) Unique one year intervention in dry season, d) Repeated interventions over five consecutive dry seasons, once a year.

Percentage of villages defined as LT hotspots in 2011 and 2012 were 35% and 31% respectively. As LT hotspots were not predictable beyond 2013, we assumed its proportion to be 31%, in order to allow forecasting. Repeating MDA and MSAT interventions in LT hotspots, once a year, during the rainy seasons, after five consecutive years, yielded a decrease in malaria incidence of 34% and 28% respectively. As interventions stopped, the efficacy reverted back and stabilized at 25%. SMC strategy reached 21% incidence decrease after 5 years delivery.

When targeting equivalent proportion of HT hotspots, repeated interventions performed equally and stabilized at 56% efficacy when delivered during the dry season. When delivered during the rainy seasons they yielded, respectively, 67% and 56% long term efficacy (Supplementary Fig. S3).

Targeting equivalent proportion of villages according to HC hotspot definitions, five years repeated interventions during the rainy seasons yielded 74% and 64% efficacy respectively for MDA and MSAT, which decreased and stabilized both at 57% at cessation of interventions. MDA and MSAT targeting HC hotspots in dry season yielded similar long-term results (Supplementary Fig. S4).

### Pre-elimination / elimination stage

Malaria elimination is defined as a decline to zero of malaria incidence in a defined geographical area as a result of deliberate efforts ^30^. Considering the continuous mathematical framework of differential equations, decline to zero incidence could not be explicitly predicted. Conversely, the elimination threshold was assessable (annual incidence below 1 case per 1,000 per year).

MDA intervention over one single year, targeting LT hotspots led to the pre-elimination stage (1-5 cases per 1,000 per year) if mosquito bites were simultaneously reduced by 10% using LLIN or Indoor Residual Spraying (IRS) throughout the year (Figure 6). The elimination stage was theoretically expectable by adding a 70% vector decrease to MDA in LT hotspots. Supplementary Table S4 summarizes the various combinations of interventions that would lead, theoretically, to the pre-elimination/elimination stage after one single year of MDA. More than a 10% simultaneous decrease in mosquito bites was needed to reach pre-elimination or elimination threshold, whatever the MDA coverage. All the above estimates took into account 80% coverage of intervention delivery and the maximum plausible weekly mobility rate (20%).

**Figure 6.**
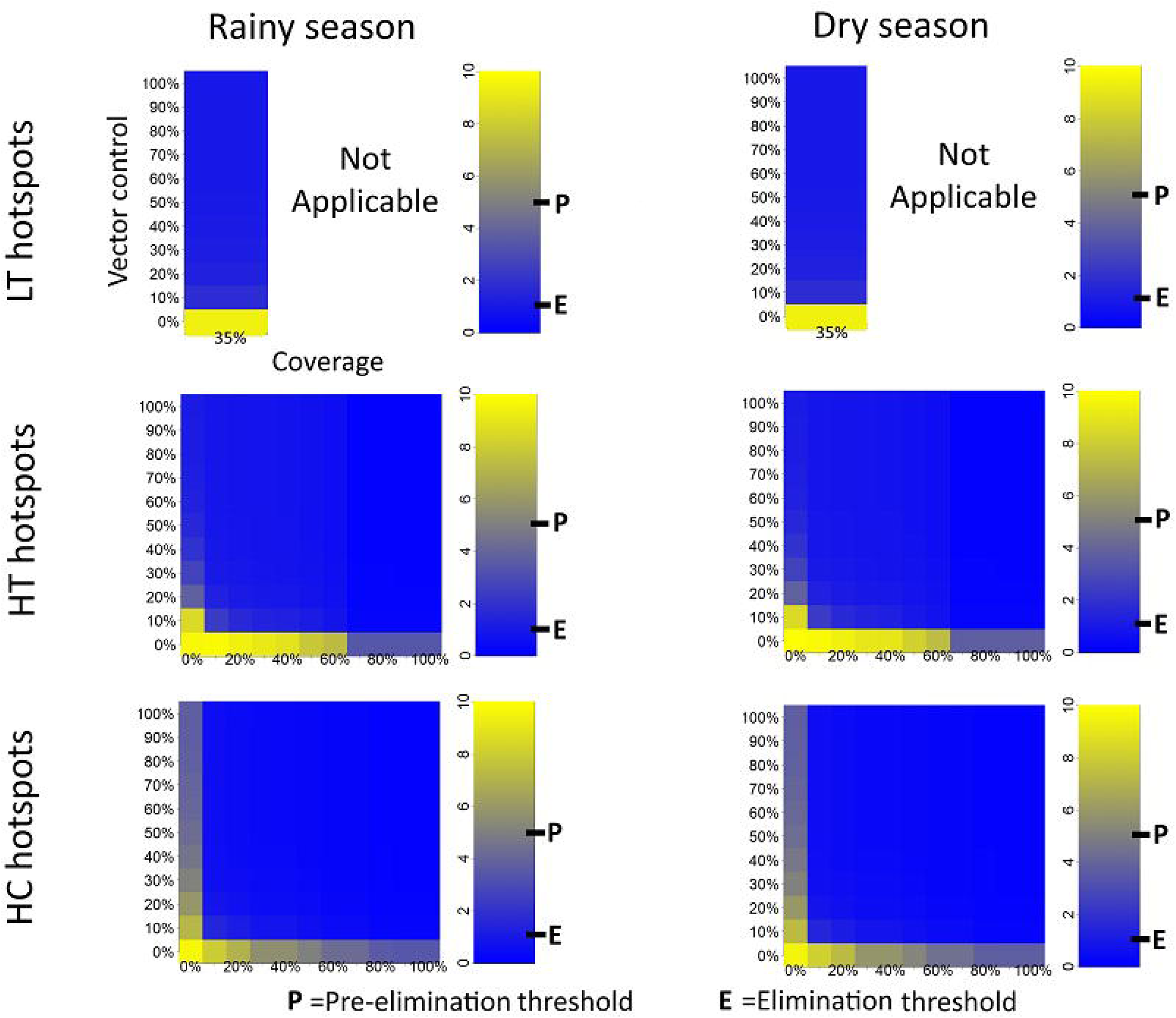
Malaria incidences in the year following simulated combined interventions (Mass Drug Administration + Vector Control). Various definitions of hotspots and different periods of intervention are tested. X-axis represents the percentage of villages included as hotspots. Y-axis represents the assumed decrease in mosquitoes bites from baseline. Color bar indicates the resulting malaria incidences and elimination/pre-elimination thresholds.

### Rebound effects due to human mobility

An incidence rebound was noticed at the cessation of repeated MDA/MSAT interventions. Rebounds occurred only under human mobility assumption. While targeting one third of HC hotspots for five consecutive rainy seasons, rebound (incidence increase) was about 17% for the overall area and 43% in targeted villages, taking into account the worst case scenario. We assumed 80% efficacy in intervention coverage, 20% asymptomatic carriage of parasites in both targeted and untargeted population and a proportion of 20% travelers in populations.

## Discussion

This study investigated the use of a spatially explicit malaria meta-population mathematical model, fitted to weekly malaria incidence in rural villages in central Senegal.

A theoretical decrease in the incidence of malaria of more than 50% was reached with both MDA and MSAT interventions repeated for five years on HT hotspots or HC hotspots (one third of villages). Reaching the pre-elimination stage (1-5 cases per 1,000 per year) was expectable only when simultaneously decreasing mosquito bites by more than 10% in MDA targeted areas. These results indicate the high impact of asymptomatic carriage and mobility on seasonal resurgence of malaria. We highlighted the foreseeable interest of spatially targeted interventions.

Obviously, the reservoir of parasites is not limited to hotspots. Our modeling framework assumed asymptomatic malaria in both targeted and untargeted areas. The asymptomatic reservoir in untargeted areas may have triggered transmission especially when mosquito bites increased at the beginning of a new rainy season. This may explain why targeting LT hotspots (15-35% of villages, supposed to be the bottleneck in dry season), was not enough to reach the elimination stage, despite the important impact of this strategy. Targeting hotspots in the dry season was intended to clear the parasite reservoir when its level was low. As widespread asymptomatic parasite carriage was assumed, high coverage repeated interventions would be needed to achieve elimination. Ideally, an optimal targeting should have been highly effective, yet with a low coverage ^8^. Asymptomatic and sub-microscopic parasite carriage should be investigated with Polymerase Chain Reaction (PCR) to display patterns of the reservoir ^31,32^. One option could be trans-sectional studies to estimate carriage prevalence ^32^. Further research is needed on the relationship between sub-microscopic parasitemia and malaria hotspot definition ^33^. It has been argued that clinical malaria incidences should not be used in hotspot definitions without taking into account asymptomatic malaria rates ^8,34^ and clustering of asexual parasite carriage using serological tools to detect malaria-specific immune responses ^8^.

Human mobility has usually been identified as a threat to malaria-free areas ^35,36^. In our study, malaria incidence decreased in untargeted areas due to the decrease in malaria importation. Some studies assumed that this incidence decrease could be related to less infected mosquitoes moving from targeted areas ^37^. Mosquito mobility modeling was not relevant in our patterns where 80% of the villages were more than 3 km far from each other.

Real human mobility data may be more accurate than radiation model. Human mobility is unlikely to be strictly related to Euclidean distances. Distance according to road network could yield more accurate analysis but this information was not available at the study area scale. Nevertheless, within the study area, the elevation is low, and relatively constant, supporting our travel assumption.

Low penetration of mobile phone and concerns about geographical scale (mobility between villages) prevented us from using Anonymized Call Details Records ^38^ to estimate mobility. Systematic studies are needed to inform mobility patterns in rural and semirural malaria areas in Senegal.

MDA interventions have contributed to eliminate malaria from islands and remote areas where population movements were closely controlled and gametocytocidal drugs have been used ^39,40^.

No resistance to Dihydroartemisinin-Primaquine was previously reported in this area and therefore not modeled. In practice, efficacy and coverage of drug interventions would also depend on the cooperation, involvement and education of local communities alongside good communication and support from local authorities ^41^.

Our meta-population mathematical model was the first to explicitly take into account human mobility at village scale, analyzing malaria transmission and interventions efficacy in Senegal.

Mathematical modeling remains an interesting tool to assess malaria strategies and policies. Nevertheless this quite deterministic approach needs to be cautiously interpreted. Unexpected changes in climatic, biological and socio-environmental factors could generate inaccuracies in predictions.

## List of abbreviations

LLIN: long-lasting insecticide-treated bednets
RDT: Rapid Diagnostic Tests
ACT: Artemisinin-based Combination Therapy
WHO: World Health Organization
MDA: Mass Drug Administration
MSAT: Mass Screen and Treat
SMC: Seasonal Malaria Chemoprevention
SEIR: Susceptible-Exposed-Infected-Recovered
GPS: Global Positioning System
MCMC: Markov Chain Monte Carlo
LT Hotspots: Low Transmission Period hotspots
HT hostpots: High Transmission Period Hotspots
HC hotspots: 3. High Connectivity Hotspots
HTP: high transmission period
LTP: low transmission period
EIR: entomological inoculation rate
IRS: Indoor Residual Spraying

## Declarations

### Ethics approval and consent to participate

Not applicable

### Consent for publication

Not applicable

### Availability of data and materials

The dataset supporting the conclusions of this article is included within the article and its additional files

### Authors’ contributions

K.S wrote the first draft of the paper. K.S, J.G and R.G developed the model. E.B and B.C coordinated data collection. M.P, R.P and K.G revised the manuscript. All authors contributed to the final manuscript.

## Acknowledgements

We thank Dr Paul Milligan for his valuable advice and for the revision of the final draft of this manuscript. This work was granted access to the HPC resources of Aix-Marseille Université financed by the project Equip@Meso (ANR-10-EQPX-29-01) of the program « Investissementsd’Avenir » supervised by the Agence Nationale pour la Recherche (Mésocentre).

---

Additional file 1

Supplementary-Information.docx : Detailed description of the mathematical meta-model

Additional file 2:

Malaria-data.xlsx: Weekly epidemiological malaria data in Mbour area

## References

1 WHO. World Malaria Report 2016, https://www.who.int/malaria/publications/world-malaria-report-2016/report/en/ (2016) (Date of access: May 11, 2017).

2 Wotodjo, A. N. et al. The implication of long-lasting insecticide-treated net use in the resurgence of malaria morbidity in a Senegal malaria endemic village in 2010-2011. Parasites & vectors 8, 267, doi:10.1186/s13071-015-0871-9 (2015).

3 Sarrassat, S., Senghor, P. & Le Hesran, J. Y. Trends in malaria morbidity following the introduction of artesunate plus amodiaquine combination in M’lomp village dispensary, south-western Senegal. Malaria journal 7, 215, doi:10.1186/1475-2875-7-215 (2008).

4 Trape, J. F. et al. The rise and fall of malaria in a West African rural community, Dielmo, Senegal, from 1990 to 2012: a 22 year longitudinal study. The Lancet. Infectious diseases 14, 476-488, doi:10.1016/S1473-3099(14)70712-1 (2014).

5 PNLP. Plan stratégique national de lutte contre le paludisme au Sénégal 2016-2020, https://www.pnlp.sn/wp-content/uploads/2016/08/PNLP_PSN_VFF_03-02-2016.pdf (2015) (Date of access: May 11, 2017).

6 Coulibaly, D. et al. Spatio-temporal analysis of malaria within a transmission season in Bandiagara, Mali. Malaria journal 12, 82, doi:10.1186/1475-2875-12-82 (2013).

7 Xu, X. et al. Microgeographic Heterogeneity of Border Malaria During Elimination Phase, Yunnan Province, China, 2011-2013. Emerg Infect Dis 22, 1363-1370, doi:10.3201/eid2208.150390 (2016).

8 Bousema, T. et al. Hitting hotspots: spatial targeting of malaria for control and elimination. PLoS medicine 9, e1001165, doi:10.1371/journal.pmed.1001165 (2012).

9 Bejon, P. et al. Stable and unstable malaria hotspots in longitudinal cohort studies in Kenya. PLoS medicine 7, e1000304, doi:10.1371/journal.pmed.1000304 (2010).

10 Bousema, T. et al. The impact of hotspot-targeted interventions on malaria transmission: study protocol for a cluster-randomized controlled trial. Trials 14, 36, doi:10.1186/1745-6215-14-36 (2013).

11 Moonasar, D. et al What will move malaria control to elimination in South Africa?, Vol. 103 (2013).

12 Prothero, R. M. Disease and mobility: a neglected factor in epidemiology. International journal of epidemiology 6, 259–267 (1977).

13 Cisse, B. et al. Effectiveness of Seasonal Malaria Chemoprevention in Children under Ten Years of Age in Senegal: A Stepped-Wedge Cluster-Randomised Trial. PLoS medicine 13, e1002175, doi:10.1371/journal.pmed.1002175 (2016).

14 R Development Core Team. R : A language and environment for statistical computing, https://www.R-project.org (2015) (Date of access: August 11, 2017).

15 Soetaert, K., Petzoldt, T. & Setzer, R. W. Solving Differential Equations in R: Package deSolve. 2010 33, 25, doi:10.18637/jss.v033.i09 (2010).

16 Soetaert, K. & Petzoldt, T. Inverse Modelling, Sensitivity and Monte Carlo Analysis in R Using Package FME. 2010 33, 28, doi:10.18637/jss.v033.i03 (2010).

17 Karney, C. F. F. Algorithms for geodesics. Journal of Geodesy 87, 43-55, doi:10.1007/s00190-012-0578-z (2013).

18 Gaudart, J. et al. Modelling malaria incidence with environmental dependency in a locality of Sudanese savannah area, Mali. Malaria journal 8, 61, doi:10.1186/1475-2875-8-61 (2009).

19 Simini, F., Gonzalez, M. C., Maritan, A. & Barabasi, A. L. A universal model for mobility and migration patterns. Nature 484, 96-100, doi:10.1038/nature10856 (2012).

20 Greenberg, J. A., DiMenna, M. A., Hanelt, B. & Hofkin, B. V. Analysis of post-blood meal flight distances in mosquitoes utilizing zoo animal blood meals. Journal of vector ecology : journal of the Society for Vector Ecology 37, 83-89, doi:10.1111/j.1948-7134.2012.00203.x (2012).

21 Doolan, D. L., Dobano, C. & Baird, J. K. Acquired immunity to malaria. Clinical microbiology reviews 22, 13-36, doi:10.1128/CMR.00025-08 (2009).

22 Ndiath, M. O. et al. Low and seasonal malaria transmission in the middle Senegal River basin: identification and characteristics of Anopheles vectors. Parasites & vectors 5, 21, doi:10.1186/1756-3305-5-21 (2012).

23 Parham, P. E. & Michael, E. Modeling the effects of weather and climate change on malaria transmission. Environmental health perspectives 118, 620-626, doi:10.1289/ehp.0901256 (2010).

24 Krefis, A. C. et al. Modeling the relationship between precipitation and malaria incidence in children from a holoendemic area in Ghana. The American journal of tropical medicine and hygiene 84, 285-291, doi:10.4269/ajtmh.2011.10-0381 (2011).

25 Doucet A, De Freitas N & Gordon N. Sequential Monte Carlo methods in practice. Vol. 1 (Springer, 2001).

26 Males, S., Gaye, O. & Garcia, A. Long-term asymptomatic carriage of Plasmodium falciparum protects from malaria attacks: A prospective study among Senegalese children. Clinical Infectious Diseases 46, 516-522, doi:Doi 10.1086/526529 (2008).

27 Jiang, B. Street hierarchies: a minority of streets account for a majority of traffic flow. International Journal of Geographical Information Science 23, 1033-1048, doi:10.1080/13658810802004648 (2009).

28 Bousema, T. et al. Identification of hot spots of malaria transmission for targeted malaria control. The Journal of infectious diseases 201, 1764-1774, doi:10.1086/652456 (2010).

29 ANSD. Situation économique et sociale du Sénégal en 2011 (French), https://www.ansd.sn/ressources/ses/chapitres/1-Demographie_2011.pdf (2011) (Date of access: March 16, 2016).

30 WHO. Malaria elimination: definitions, criteria and possible variants, https://www.who.int/malaria/mpac/malaria_elimination_definitions_criteria_presentation.pdf (2013) (Date of access: June 30, 2016).

31 Imwong, M. et al. The epidemiology of subclinical malaria infections in South-East Asia: findings from cross-sectional surveys in Thailand-Myanmar border areas, Cambodia, and Vietnam. Malaria journal 14, 381, doi:10.1186/s12936-015-0906-x (2015).

32 Gerardin, J. et al. Optimal Population-Level Infection Detection Strategies for Malaria Control and Elimination in a Spatial Model of Malaria Transmission. PLoS Comput Biol 12, e1004707, doi:10.1371/journal.pcbi.1004707 (2016).

33 Kangoye, D. T. et al. Malaria hotspots defined by clinical malaria, asymptomatic carriage, PCR and vector numbers in a low transmission area on the Kenyan Coast. Malaria journal 15, 213, doi:10.1186/s12936-016-1260-3 (2016).

34 Sturrock, H. J. et al. Targeting asymptomatic malaria infections: active surveillance in control and elimination. PLoS medicine 10, e1001467, doi:10.1371/journal.pmed.1001467 (2013).

35 Silal, S. P., Little, F., Barnes, K. I. & White, L. J. Hitting a Moving Target: A Model for Malaria Elimination in the Presence of Population Movement. PloS one 10, e0144990, doi:10.1371/journal.pone.0144990 (2015).

36 Silal, S. P., Little, F., Barnes, K. I. & White, L. J. Towards malaria elimination in Mpumalanga, South Africa: a population-level mathematical modelling approach. Malaria journal 13, 297, doi:10.1186/1475-2875-13-297 (2014).

37 Bousema, T. et al. The Impact of Hotspot-Targeted Interventions on Malaria Transmission in Rachuonyo South District in the Western Kenyan Highlands: A Cluster-Randomized Controlled Trial. PLoS medicine 13, e1001993, doi:10.1371/journal.pmed.1001993 (2016).

38 Bengtsson, L. et al. Using mobile phone data to predict the spatial spread of cholera. Scientific reports 5, 8923, doi:10.1038/srep08923 (2015).

39 Kaneko, A. et al. Malaria eradication on islands. Lancet 356, 1560-1564, doi:10.1016/S0140-6736(00)03127-5 (2000).

40 Song, J. et al. Rapid and effective malaria control in Cambodia through mass administration of artemisinin-piperaquine. Malaria journal 9, 57, doi:10.1186/1475-2875-9-57 (2010).

41 Kondrashin, A. et al. Mass primaquine treatment to eliminate vivax malaria: lessons from the past. Malaria journal 13, 51, doi:10.1186/1475-2875-13-51 (2014).

